# Efficient storage and analysis of quantitative genomics data with the Dense Depth Data Dump (D4) format and d4tools

**DOI:** 10.1101/2020.10.23.352567

**Authors:** Hao Hou, Brent Pedersen, Aaron Quinlan

## Abstract

Modern DNA sequencing is used as a readout for diverse assays, with the count of aligned sequences, or “read depth”, serving as the quantitative signal for many underlying cellular phenomena. Despite wide use and thousands of datasets, existing formats used for the storage and analysis of read depths are limited with respect to both file size and analysis speed. For example, it is faster to recalculate sequencing depth from an alignment file than it is to analyze the text output from that calculation. We sought to improve on existing formats such as BigWig and compressed BED files by creating the Dense Depth Data Dump (D4) format and tool suite. The D4 format is adaptive in that it profiles a random sample of aligned sequence depth from the input BAM or CRAM file to determine an optimal encoding that often affords reductions in file size, while also enabling fast data access. We show that D4 uses less storage for both RNA-Seq and whole-genome sequencing and offers 3 to 440-fold speed improvements over existing formats for random access, aggregation and summarization. This performance enables scalable downstream analyses that would be otherwise difficult. The D4 tool suite (d4tools) is freely available under an MIT license at: https://github.com/38/d4-format.

## Introduction

Aligned DNA or cDNA sequence depth is one of the most important quantitative metrics used for variant detection^1^, differential gene expression^2,3^, and for critical evaluations of data quality control^4^. Despite the wide use of quantitative genomics datasets, the underlying algorithms, data structures, and software implementations for handling genomics quantitative data have limitations. The BigWig format^5^, a workhorse in genomics, requires considerable memory during its creation and is complex enough to limit broad development of, or improvements to, the format. At the other end of the spectrum is the widely-used, BEDGRAPH format which, owing to the fact that it is text-based, is simple to understand, yet both slow to parse and consumes substantial disk space.

To improve upon the limitations of existing formats, we have developed the Dense Depth Data Dump (D4) format and software suite. The D4 format is motivated by the observation that depth values often have low variability and are therefore highly compressible. Here, we detail how we use this low entropy to efficiently encode quantitative genomics data in the D4 file format. We then demonstrate the D4 format’s combined file size and analysis speed efficiency with respect to the bgzipped BEDGRAPH, BigWig^5^, and HDF5^6^ formats. The D4 format and associated tools support fast random access, aggregation, summarization, and extensibility to future applications. These capabilities facilitate a new scale of genomic analyses that would be otherwise far slower.

## Methods

### The D4 format

Devising a disk- and computation-efficient file format to store quantitative data that can take on an infinite or very large range of finite values is a difficult task. Thankfully, the sequencing depths observed in modern genomics assays often have little variance, thus yielding a limited range of discrete values. For example, consider a human whole genome sequence (WGS) assay yielding the typical target of 30-fold average read depth. In a typical WGS dataset, more than 99% of the observed depths fall between 0 and 63 (**Figure 1A**). Therefore, it is possible to encode >99% of the data using only 6 bits per base, since 2^6^ equals 64. Similarly, more than 50% of genomic positions have a sequence depth of 0 in typical RNA-seq experiments, since a small portion of the coding genome is assayed. On the other hand, RNA-seq yields a much broader range of non-zero depths than WGS, reflecting the highly-variable degree of isoform expression from gene to gene. Nonetheless, the range of observed values is both finite and redundant (**Figure 1B**).

**Figure 1.**
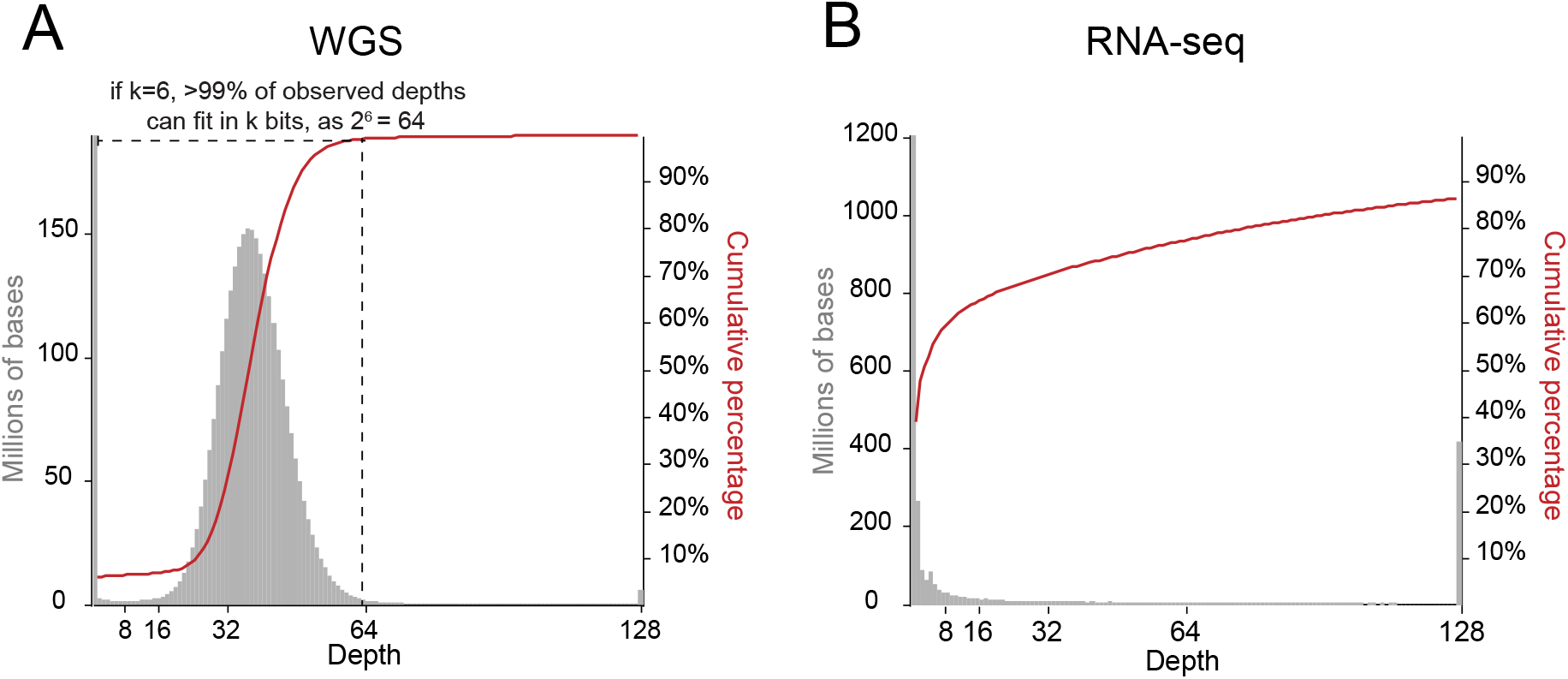
Depth distribution for WGS and RNA-seq datasets. The global depth histogram (gray) and cumulative percentage of bases (red) for whole-genome (**A**) and RNA-Seq (**B**) data. For typical WGS datasets, more than 99% of observed depth values are <63, indicating that the majority of data can be encoded in 6 bits. The variance in observed depths is much greater for RNA-seq data and other quantitative assays; however, the range of values is finite and amenable to encoding. The WGS dataset is sample HG002 from the Genome in a Bottle Project, and the RNA-seq dataset is sample ENCFF976QSN from the ENCODE project^7^.

The D4 format utilizes an encoding scheme that takes advantage of the finite range of depth values observed in most genomics assays to enable storage efficiency (**Figure 2**). D4 uses a dense array as a primary table having one entry per base in a given chromosome with each base consuming *k* bits (**Figure 2A**, where *k*=6). The index *i* into this dense array provides a lookup for the chromosomal position *i*. That lookup is then used as a key into a *k*-bits code table mapping 2^k^ *k*-bit codes to the 2^k^ distinct real values (**Figure 2B**). We note that the Code Table could be used to encode other values besides sequencing depths (e.g. labels, p-values, etc.), but the number of distinct codes in the Code Table is governed by the choice of *k*. Any value less than 2^k^-1 in the code table encodes the actual depth for position *i*. However, if the encoded value is exactly 2^k^-1, then either the value is encoded by the code 2^k^-1, or its precise value is stored in the sparse secondary table (**Figure 2C**). Therefore, when a code 2^k^-1 is seen, the sparse secondary table must be queried. If this query of the secondary table does not return an entry for that position, one can conclude that the actual value is the value encoded by the code 2^k^-1. Otherwise the actual value is returned from the secondary query. In effect, if a value cannot be encoded to a *k*-bit code by the code table, the code 2^k^-1 is emitted to the primary table as a placeholder, and the actual value is stored in the sparse secondary table instead.

**Figure 2.**
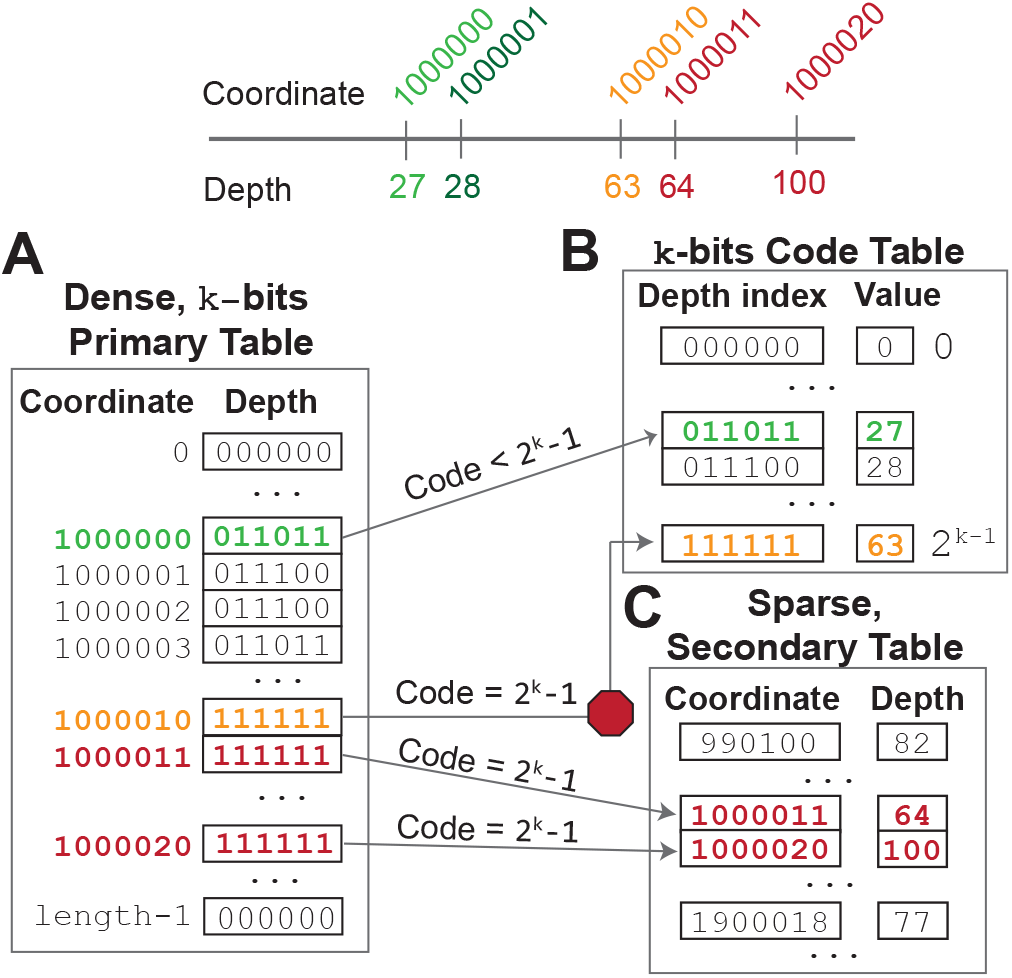
The D4 format encoding strategy. A. The D4 Dense Primary Table. The D4 format uses a dense array as a Primary Table that contains one entry per base in each chromosome. Each array entry consumes k bits, and the values stored in each entry range from 0 to 2^k^ - 1. In this hypothetical example, k is 6. **B. The k-bits Code Table**. After a D4 file is created and when one wants to learn the depth of coverage at a particular genome position, one looks at the code stored at the position in the primary table. If the value of the code is less than 2^k^ - 1, then one can look up the actual value (e.g., sequencing depth) in the k-bits Code Table. For example, the code for position 1,000,000 is “011011”, which is less than 2^6^ − 1 (63). Therefore, the code table is used to look up the encoding for “011011”, which, in this case, is 27. **C. The Sparse Secondary Table**. There is more work to be done in cases where the code stored in the Primary Table is exactly equal to 2^k^ - 1. This scenario indicates that either the value for that position is encoded in the last possible entry of the code table, or that the value exceeds the range of distinct values that can be encoded by the choice of k. To distinguish the two scenarios, one must first lookup the genome coordinate in the sparse Secondary Table. If, as in the case of coordinate 1,000,010, an entry does not exist in the Secondary Table (denoted by the red “stop sign”), then we can infer that the actual value can be determined by looking up the code in the Code Table. If, however, an entry does exist, as in the cases of coordinate 1,000,010 (depth = 64, which exceeds the range of 0 to 2^k^-1) and 1,000,020 (depth = 100), then the value for that coordinate is stored directly in the Secondary Table.

For a typical 30X WGS, this secondary table query happens less than 1% of the time and therefore the sparse secondary table often contains less than 1% of the data. This efficiency saves disk space as the primary table uses *k* bits per base, and the secondary table usually contains a small percentage of the data but uses 80 bits to store the combination of the chromosomal range and the observed depth at that range of positions. In this way, we estimate that chromosome 1 of the human genome, which contains 249 million bases, will consume 212 megabytes for a D4 encoding of WGS data, as 249 million * 6 bits + 1% * 249 million * 80 bits < 181 megabytes. In contrast, if we were to store each depth as an unsigned 32-bit integer, 996 megabytes would be required for the same data. Therefore, we can achieve a 5.2X lossless encoding ratio with the D4 encoding strategy with *k* equal to 6. Usually for WGS datasets, the secondary table encodes only a small portion of the values, but for RNA-seq datasets the optimal k is 0, which means we have a zero-sized primary table and all the data is completely stored in the secondary table (discussed further below). In order to further reduce file size, D4 allows an optional compression for the secondary table. When this is enabled, all the secondary table records are compressed in blocks which usually reduces the secondary table size by more than 50% for RNA-seq datasets.

### Adaptive Encoding

The toy example in **Figure 2** assumes *k* is known ahead of time. However, for each dataset, there is an unknown, optimal *k* which maximizes the efficiency of the read depth encoding. This optimal *k* depends on the range and variance of depths observed in the dataset. The d4tools software samples the depth distribution to quickly determine the optimal *k* for a given dataset. The essential trade-off is that a smaller *k* necessarily means that each entry in the primary array will use fewer bits (**Figure 3**). However, with a smaller *k*, a greater percentage of the observed depths will fall outside of the range supported by the code table given the choice of *k* (e.g., depths 0 through 63 are supported by *k*=6). Likewise, a larger *k* means that each base in the primary array will consume more bits, but that fewer values are needed in the sparse secondary table, which consumes 80 bits per entry. As mentioned, in the case of most 30X WGS datasets the optimal choice of *k* is 6 (**Figure 3A**). However, the optimal choice of *k* for most RNA-seq datasets is 0. This is because the range of depths observed in RNA-seq data is wide enough that it’s not possible to choose a narrow range (small *k*) that encompasses the majority of the observed depths (**Figure 3B**). When *k* becomes too large, each entry in the chromosome-length primary table will consume enough bits to offset any gain from the larger range.

**Figure 3.**
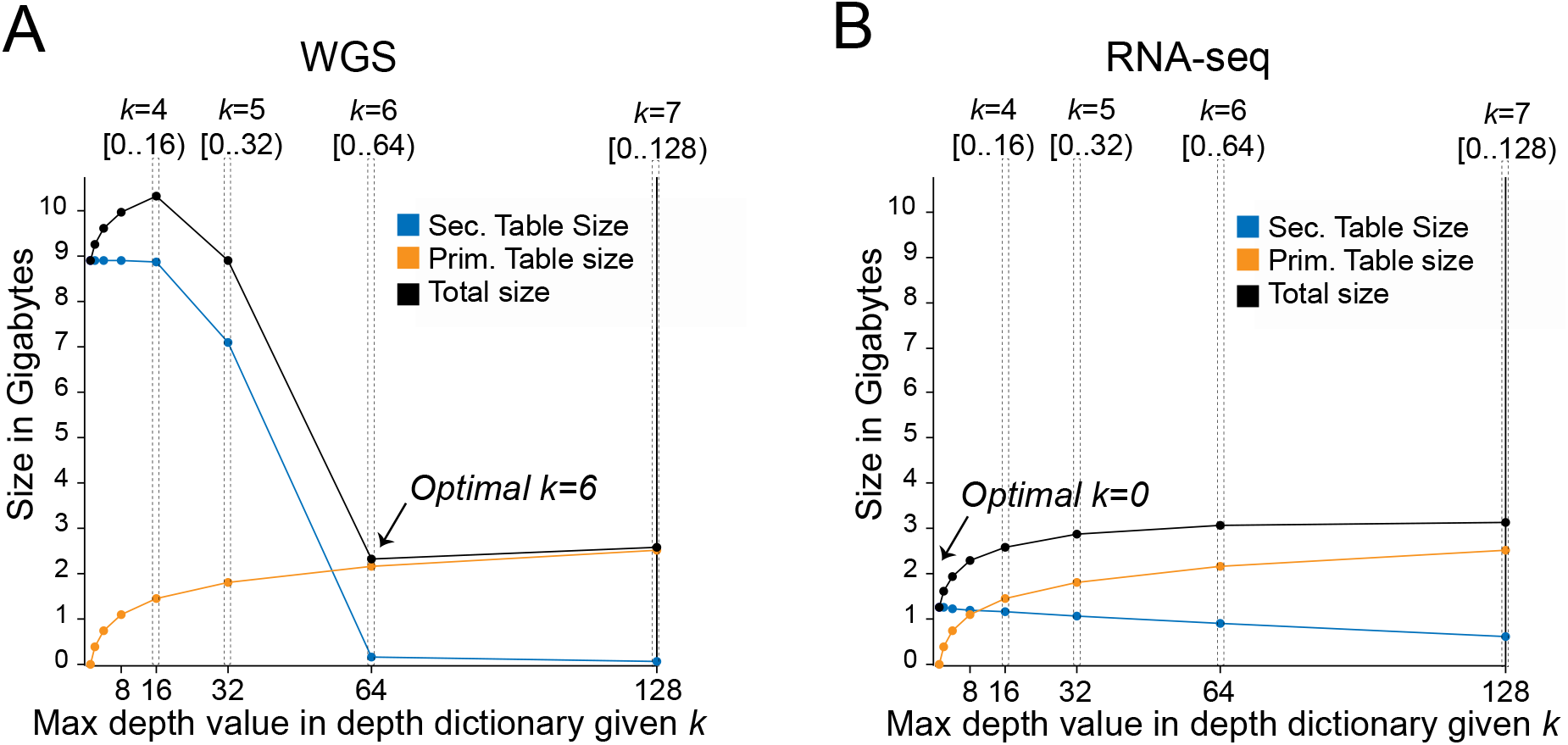
Optimizing the choice of *k* given the trade-off between the size of the primary and secondary tables. Panels A and B demonstrate, for example WGS and RNA-seq datasets, respectively, the trade-off between the size of the primary and secondary tables *k* is varied. The optimal choice of *k*, which minimizes the total size of the resulting D4 file, is the point at which the black line has its lowest value. Each entry in the primary table will consume more bits as *k* increases, but the secondary table size will decrease. D4 finds the optimal *k* by randomly sampling the depth distribution from the input file. Note that for RNA-Seq (panel B), it is often optimal to have *k* of zero, indicating that only the sparse secondary table is used.

### Enabling parallelism, modularity, and extensibility in the D4 format

The implementation of the D4 format uses a general data container that allows multiple, variable-length data streams to be appended to the container file in parallel (**Supplementary Methods**). In particular, when a D4 file is being created, genome coordinates are split into small chunks. Each thread is responsible for several chunks of data, encoding and writing the data to the growing D4 file in parallel. The container file data structure is based on an unrolled list^8^ with adaptive chunk sizes. When a D4 file is created, it uses the unroll list to allow data chunks to be added to the growing D4 file without blocking the other threads. When reading a D4 file, the unrolled list can be mapped to the main memory using the “mmap” system call and manipulated with the CPU’s SIMD instruction set. D4 files are able to be extended without breaking any existing file format by adding more streams to the container file, for example, a precomputed depth statistic.

### An efficient algorithm for computing depth of coverage

In order to fully take advantage of the encoding efficiency of the D4 format, we developed a new algorithm to efficiently profile the depth of coverage in input BAM files. We developed this algorithm because existing methods such as samtools^9^ “pileup” consume memory in proportion to the maximum depth in a region. This approach is also slow in genomic regions exhibiting very high depth. The memory use for mosdepth^10^, our previously published method for reporting per-base sequencing depth, is governed by the length of the longest chromosome, but, as implemented, it is not parallelized. In contrast, D4 introduces a new algorithm that limits the memory dependency on depth and also facilitates parallelization of the coverage calculation. This algorithm uses a binary heap that fills with incoming alignments as it reports depth. The average time complexity of this algorithm is linear with respect to the number of alignments (see **Supplementary Methods** for algorithm details and lower bound analysis). Because there is little memory allocation, this algorithm can be parallelized by performing concurrent region queries in different threads. The algorithm uses the start and end of alignments such that it does not count, for example, soft-clipped regions, but it will also not drop the depth for internal CIGAR events such as deletions. As a result, the d4tools “create” command is able to encode new D4 files from input BAM or CRAM files in far less time than creating depth profiles with BigWig or other formats, especially when leveraging multiple threads.

### The d4tools software

The d4tools software suite is written in the Rust programming language to facilitate safe concurrency and performance. Our implementation takes advantage of Rust’s trait system, which allows zero-cost abstraction and the borrow checker to ensure correctness of the memory management. More importantly, Rust provides a strong multithreading safety guarantee which allows us to implement D4 in a highly parallelized fashion. We expose a C-API and provide a separate Python API so that researchers can easily utilize the library. Example commands for common operations to create and analyze D4 files are provided in **Table 1**.

**Table 1.** Example d4tools commands.

| Command | Description |
| --- | --- |
| d4tools create input.bam output.d4 | Create a d4 file by profiling depth-data from an alignment file |
| d4tools create input.bigwig output.d4 | Create a d4 file by profiling depth-data from a BigWig file |
| d4tools create -g input.genome input.bedgraph output.d4 | Create a d4 file by profiling depth-data from a BedGraph file |
| d4tools view output.d4 chr1:1-100000 | Extract data for a given region to text-based BED format |
| d4tools stat -s mean -r regions.bed output.d4 | Calculate the mean coverage for each region given in a BED file |
| d4tools plot output.d4 chr1:1000-2000 | Draw an image of coverage data for the given region. |

## Results

### Single-Sample Evaluation

In order to compare D4 to existing solutions, we calculated aligned sequence depth profiles for the WGS and RNA-seq samples in **Figure 1** using the D4, BGZF-compressed BedGraph, BigWig, and HDF5 formats. In addition to comparing the file sizes for each approach, we evaluated the time required to create a depth profile from each aligned BAM file into the relevant file format, the time required to summarize the results in a full sequential scan of the entire file, and finally, the average time used to query the depth for a set of random genomic intervals (**Figure 4**). All operations by D4 yield an increase in performance as the number of threads increases, although this performance increase begins to saturate for random access around 8 cores (**Supplementary Figure 1**). For both WGS and RNA-seq datasets, D4 yielded a 10X faster file creation time, and, with the exception of the highly-compressed HDF5 format, yielded the smallest file size. The high compression rate of the HDF5 format comes at the cost of much less efficient analysis times. Sequential access to the depth at every genomic position was 3.6 times faster than BigWig, 3.8 times faster than HDF5 with a single thread and 31.5 and 32.4 times faster with 64 threads. Again using 1 and 64 threads, accessing 10,000 random genomic intervals is between 21.3 and 72.8 times faster than BigWig, between 130 and 446 times faster than block-GZIPed BEDGRAPH and between 18 and 64 times faster than HDF5. D4 also supports optional secondary table compression. For WGS datasets, the secondary compression usually doesn’t result in a noticeable performance and size difference, but for RNA-seq datasets, enabling the secondary table compression reduces the file size at the cost of slower computation. Comparing a D4 file with the deflated and inflated secondary table, the D4 file is 54% smaller while the sequential access performance of a deflated D4 file is 4 times slower with single thread and 44% slower with 64 threads. Similarly, the random-access performance is slower by 17 times and 2 times. In practice, D4 files with deflated secondary tables can be used as a compact archive format for cold data and those with inflated secondary tables can be used as hot data format that allows high performance data analysis.

**Figure 4.**
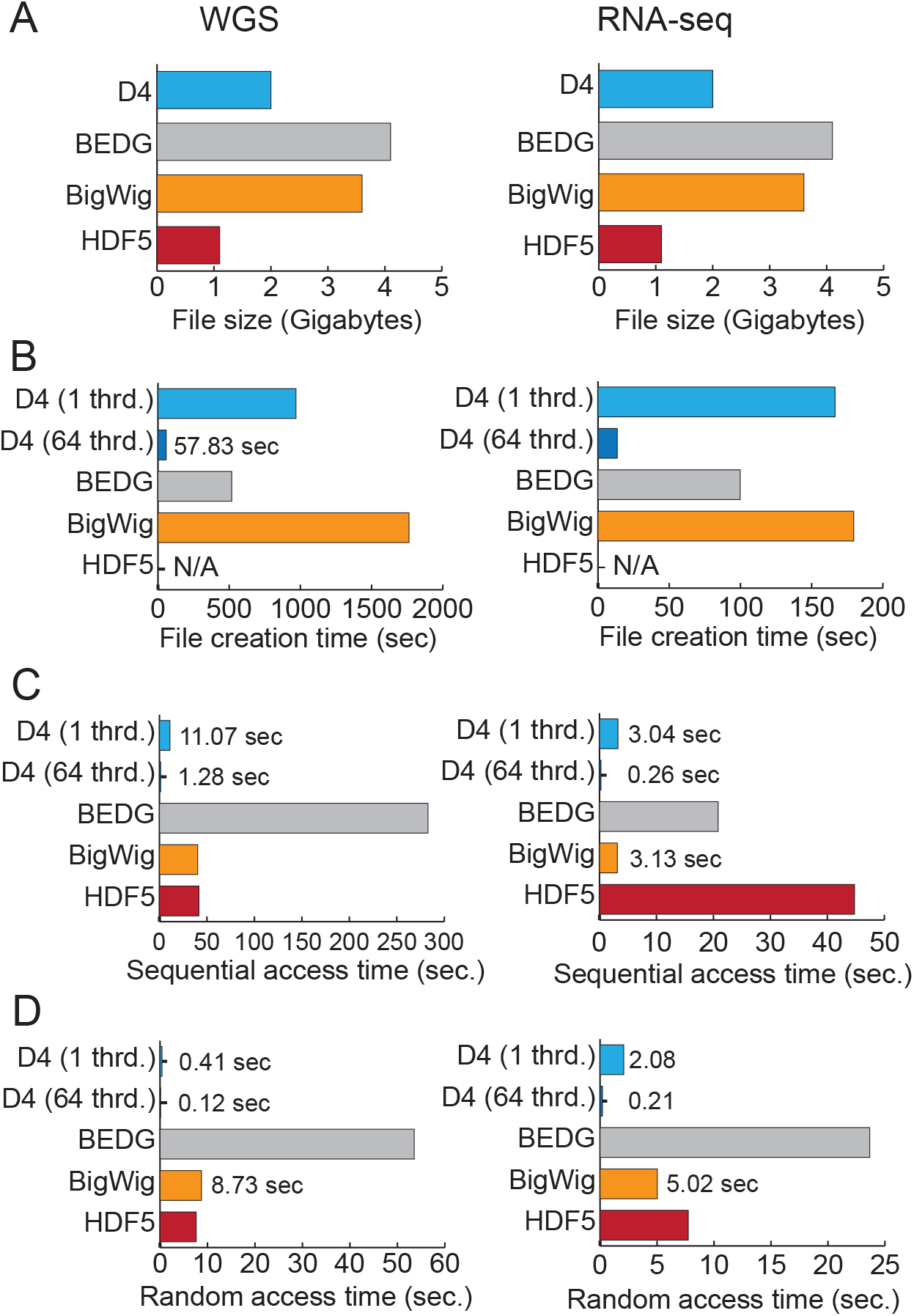
Performance of D4 compared to other formats. The file size (**A**), total wall time for creation (**B**), sequential access (**C**) and random access (**D**) are reported for the WGS and RNA-Seq datasets presented in Figure 1. The “sequential access” experiment iterates over each depth observed in the output file. Ten thousand random intervals of length 10,000 were used to assess the “Random Access” time. Each time reflects the average of 5 trials with the min and max removed. All D4 files shown use a compressed secondary table. Thrd: abbreviation for threads.

### Evaluation on large cohort

In order to illustrate the types of analyses facilitated by the speed and efficiency offered in D4 format and tools, we performed an evaluation of depth on both the 2,504 samples from 1000 Genomes high-coverage whole-genome samples (Michael Zody, personal communication), and on 426 RNA-seq BAM files from the ENCODE project. We restricted our comparison to the BigWig format given its wide use in the genomics community, and the fact that its combined file size and performance are closest to that of D4. For WGS datasets, D4 files are consistently less than half the size of BigWig files and D4 file size is largely consistent across a wide range of input CRAM file sizes (**Figure 5A**). As the mean depth of WGS datasets from the 1000 Genomes project increases, we observe a transition from 5 to 6 in the optimal choice of k, since the variance increases with the mean (**Figure 5A**, inset). This results in a small increase in D4 file size, yet we observed a large jump in BigWig file size for WGS datasets with a mean depth near 60 or higher. Furthermore, we note that the male 1000 Genomes datasets had slightly larger D4 files than females owing to the fact that one X chromosome yields more positions with aligned sequencing depths that lie outside of the range encoded by *k*. Finally, while D4 files are much smaller than BigWig for WGS datasets, D4 and BigWig file sizes are much more similar for RNA-seq datasets (**Figure 5B**).

**Figure 5.**
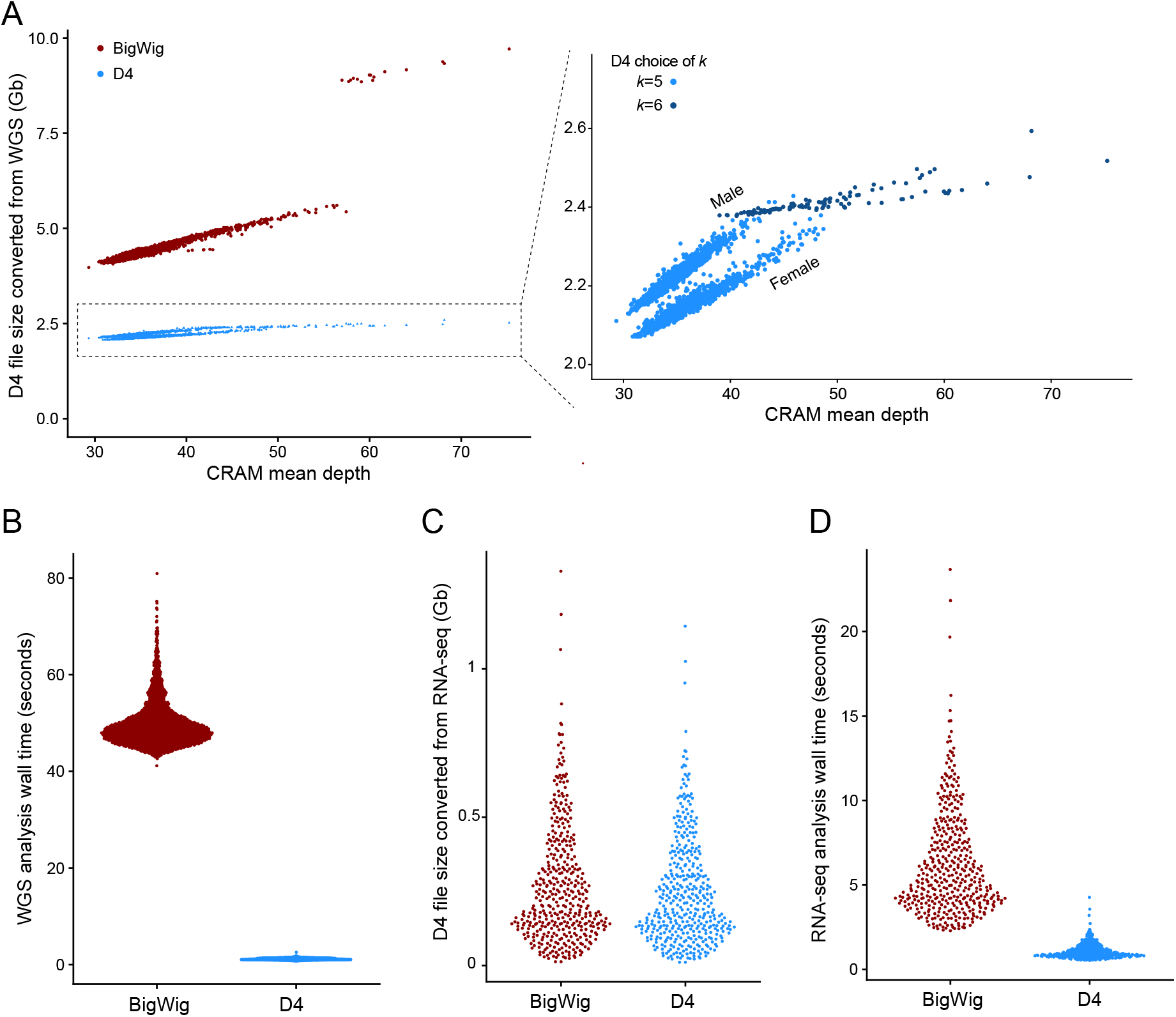
**A**. Mean depth of the original alignment file (x-axis) compared to the size of the resulting D4 and BigWig files (y-axis). Note the small range of D4 file sizes. The inset figure depicts how the optimal choice of *k* changes as mean depth increases. Furthermore, male samples have slightly higher D4 file sizes than female (see main text). **B**. Size distribution of the D4 and BigWig files converted from RNA-seq BAM files. **C**. Per-sample distribution of wall time required to identify regions of each genome greater than or equal to 1,000 base pairs where each position has a depth of coverage greater than 120. 98.8% of D4 samples completed in under 2 seconds of wall time each. **D**. Per-sample distribution of wall time required to identify regions of each genome greater than or equal to 1,000 base pairs where each position has a depth of coverage greater than 10,000.

As an example of the type of practical analyses improved by use of D4, we identify large regions in the WGS and RNA-seq datasets where samples had high coverage depths, as these often reflect atypically high depth regions that result in incorrect variant calls^11^, and genes with high expression levels, respectively (**Figure 5C,D**). We implemented this using the D4 API in Rust, and the relevant code can be found in the D4 Github repository. We report the time to create each D4 file and the time to calculate the high-coverage regions. On average, D4 required only 3% of the time required by BigWig to conduct these analyses for the WGS datasets, and 18% of the time required by BigWig for the RNA-seq datasets.

## Discussion

We have introduced the D4 format and set of tools and shown the speed, scalability, and utility. Our results illustrate D4’s unique combination of minimal file size and computational efficiency in addition to highlighting the potential of D4 for rapid analysis of large-scale datasets. In particular, unlike existing formats, the adaptive encoding strategy in D4 allows the format to adjust to the properties of diverse genomics datasets having varied data densities such as whole-genome sequencing, RNA-seq, ChIP-seq, and ATAC-seq.

Looking forward, we emphasize that the flexible architecture of the D4 format will allow it to adapt to other applications such as representing signals from single-cell RNA sequencing datasets where multiple layers of information (e.g., cell ID, barcode, etc.) are required.

Furthermore, the D4 format’s efficiency can also facilitate rapid data visualization for genome browsers, as well as rapid statistical analyses and dataset similarity metrics that leverage additional indices or precomputed metrics stored in the D4 format. For example, we have already incorporated D4 output into our popular depth-profiling tool, mosdepth^10^. This change resulted in a nearly 3-fold reduction in run time, since much of the original run time was spent on writing output in compressed BEDGRAPH format. In addition, the D4 output from mosdepth is more amenable to custom downstream analyses.

Given its speed, flexibility, and the associated Rust and Python programming interfaces, we anticipate that the D4 format will enable large-scale genomic analyses that remain challenging or intractable with existing formats.

## Code availability

D4utils: https://github.com/38/d4-format

D4 Rust API: https://docs.rs/d4

D4 Python API: https://github.com/38/pyd4

## Acknowledgements

We acknowledge helpful comments from members of the Quinlan laboratory, as well as funding from the National Institutes of Health https://docs.rs/d4-framefile(NIH) grants HG006693, HG009141 and GM124355 awarded to A.R.Q.

## SUPPLEMENTARY FIGURES

**Supp Figure 1.**
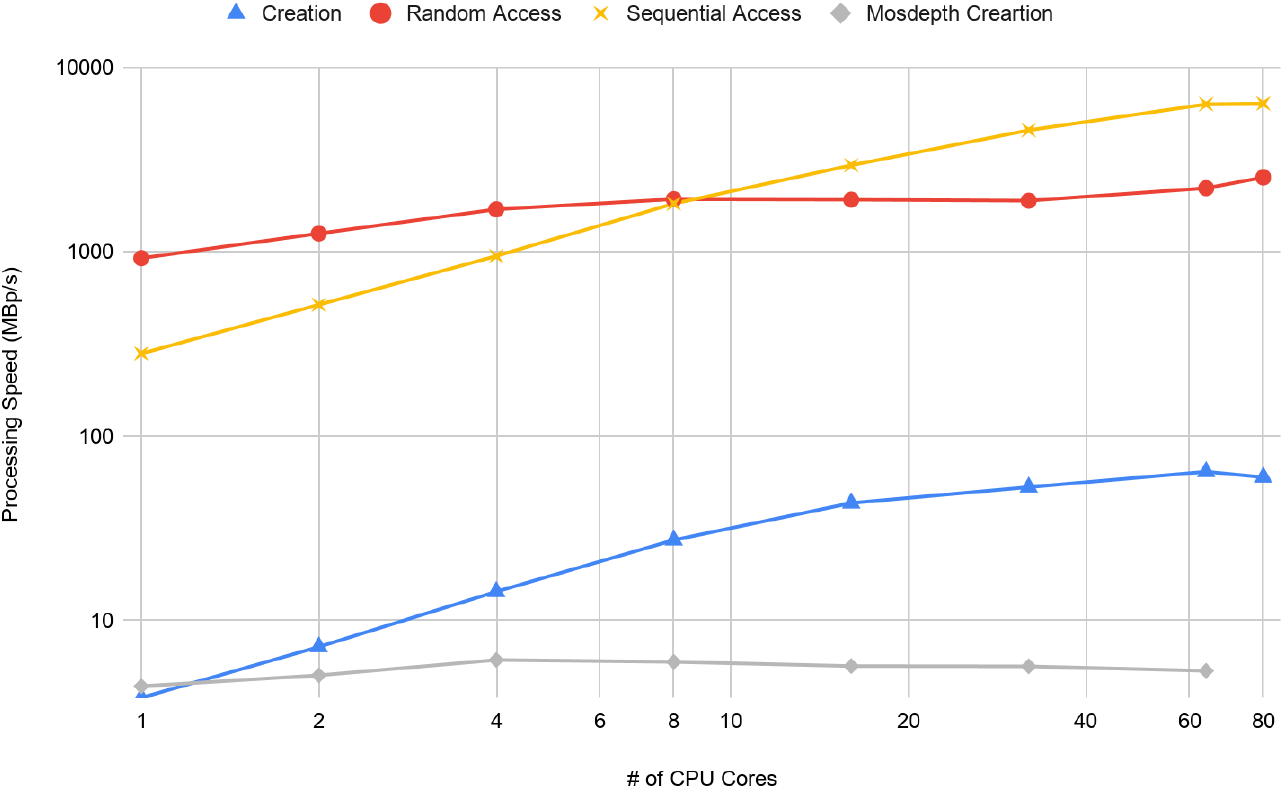

## SUPPLEMENTARY METHODS

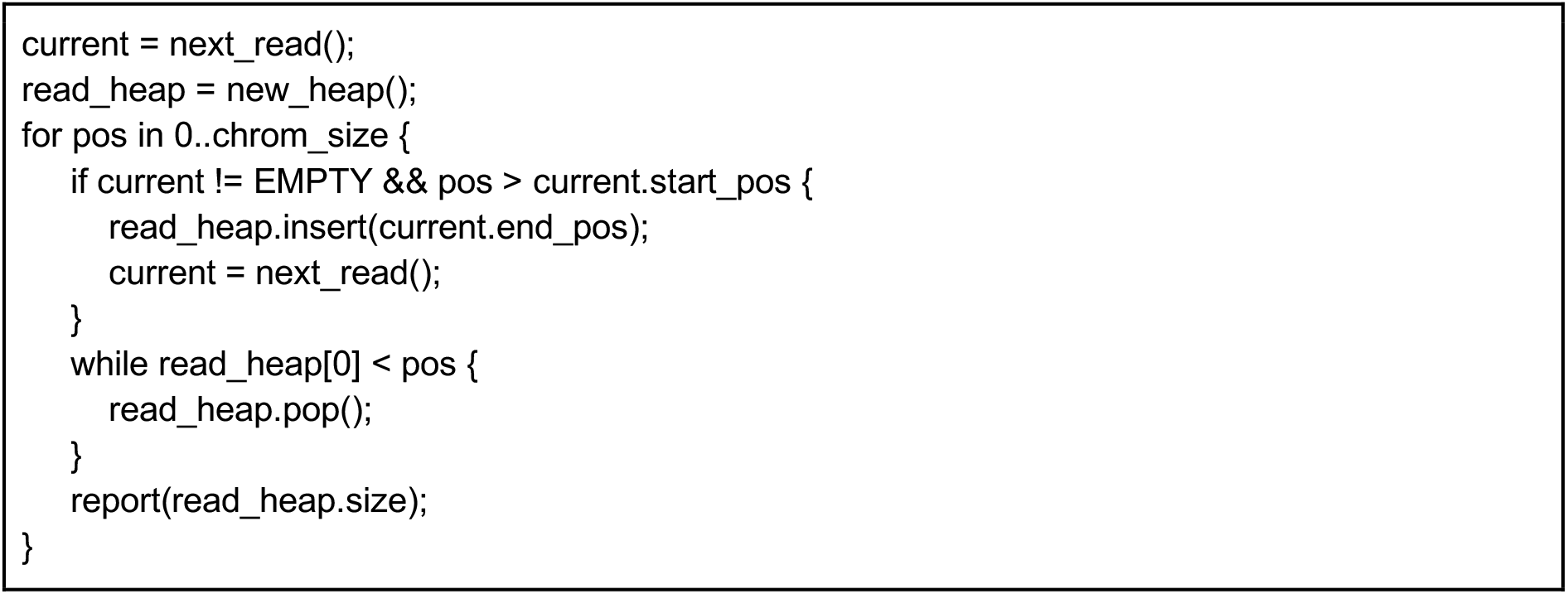
Depth profiling algorithm.

For each iteration, the binary heap operation takes O(log(D)) running time where D is the current read depth, thus, we can prove that:

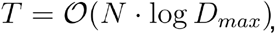

where N is the genome size and D_max_ is the maximum depth. Now we prove a tighter bound for it.

For the theoretical lower bound, we should at least enumerate all the reads in the alignment file. Because we need to both scan the alignment file and report the per-base values even in the worst case. So that

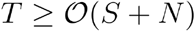

where S is the total number of reads in the alignment file. The accurate running time of our algorithm is

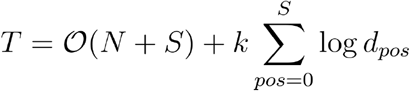

where d_pos_ is the depth at position pos. Applying the inequality of arithmetic and geometric means, we can easily show that

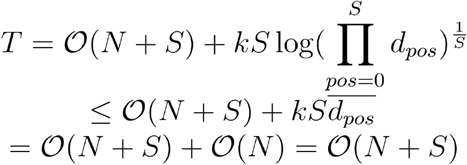

Which means our depth profiling algorithms is no worse than the theoretical upper bound, so that

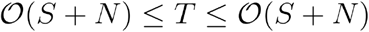

So that we proved that

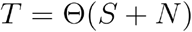

It’s possible to (1) simplify the alignment in the read_heap to a single end position and (2) collapse all the alignment ends at the same location. By doing so, the upper bound of memory efficiency is

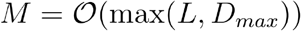

where L is the maximum of the read length.

So that this algorithm is optimal in terms of time complexity while space complexity is better than the counter array approach. Thus, this algorithm allows the BAM file to be processed with multiple processors at the same time with limited memory usage.

### D4 file format details

#### Overview

Unlike normal file formats, D4 doesn’t have a stable file layout. The D4 file is built on top of the container format called framefile. All the data including primary table, secondary table and metadata is stored as an entity of the framefile.

There are 3 different types of object in a framefile: variant length data stream, fixed-sized blob and sub-framefile. In the D4 implementation, the primary table is implemented as a fixed-sized blob and the secondary table is implemented as a sub-framefile that contains multiple data streams of sorted-by-position out-of-range sparse values.

Unlike the traditional file IO API, our D4 implementation doesn’t use read/write semantics to perform the IO. When a D4 file is open, the D4 file can be split into small chunks (e.g. 1,000,000,000 bps per chunk), each chunk can be handled by different threads at the same time. For random access cases, the irrelevant chunks are dropped once the file has been split.

#### Splitting the Primary Table

For both read and write, the primary table is directly mmap-ed to the address space of the program and it’s directly split into slices.

#### Splitting the Secondary Table

For writing a D4 file, each thread will create an independent stream under the secondary sub-framefile and each of the streams is called a partition of the secondary table. When a D4 file is being read, all the frames will be indexed into memory first and then splitted into small pieces according to the primary table partition.

#### D4 File Format Overview

The D4 format is defined as a file header and the root container

| Offset (in bytes) | Name | Type | Value |
| --- | --- | --- | --- |
| 0 | File Magic Number | [u8;4] | "\xdd\xddd4" |
| 4 | Format Version | [u8;4] | [0,0,0,0] |
| 8 | Frame File Root | Directory | Primary Size = 512 |
| Frame File Data |  |  |  |

### Frame File Structure

#### Streams

A Stream is a variant-length series of bytes. In framefile container format, a stream is defined as a linked list of frames:

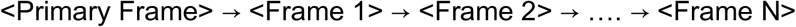

The first frame is called the primary frame.

#### Stream Frame

Represents a single part of the stream, the format is defined as the following table, the offset is relative to the start of the frame. (Physical Layout)

| Offset (in bytes) | Name | Type | Note |
| --- | --- | --- | --- |
| 0..4 | next_frame | u32_le | Relative Offset in Bytes (For the last frame, must be 0) |
| 4..8 | next_size | u32_le | Data Chunk size of next frame in bytes |
| 8..8+size | Data Chunk |  |  |

#### Directory

Dictionary is a special form of stream, data payload is a list of directory entries and list is terminated with a byte 0. (Logic Layout)

|  |
| --- |
| Entry 1 |
| Entry 2 |
| ... |
| Entry N |
| '\0' |

#### Directory Entry

(Logic Layout)

Directory Entry (Logic Layout)
| Offset (bytes) | Name | Type | Note |
| --- | --- | --- | --- |
| 0 | valid_flag | u8 | Must be 1 |
| 1 | entry_type | u8 | 0 = Stream<br>1 = Sub-framefile<br>2 = Fix-size blob |
| 2..10 | primary_frame | u64_le | Absolute offset to the primary frame for this entity |
| 10..18 | primary_size | u64_le | Size of the primary frame<sup>12</sup> |
| 18..? | name | null terminated string | Name of the entity (UTF8 encoding) |

1. For Sub-directory, this is defined as size of everything under the sub dir 2. For blob, it defined as blob size

#### Blob

Any binary data with known size

#### D4 Logical Representation Overview

D4 file is defined on the top of the container format. The d4utils provides a tool to inspect the logic structure of a D4 file with “d4utils framedump”

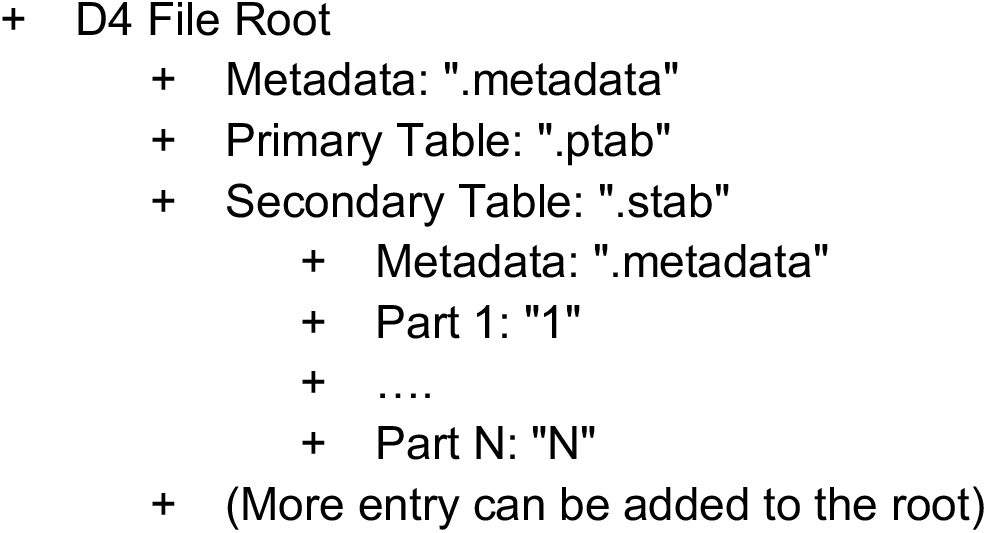

| Name | Path | Type | Description |
| --- | --- | --- | --- |
| Metadata | .metadata | Stream | Json encoded global metadata, e.g.<br>{<br>"chrom_list": [ |
|  |  |  | `{ "name":"chr1", "size": 10000000 } ], "dictionary": { "simple_range": { "low": 0, "high": 64, } } }` |
| Primary Table | .ptab | Blob | K-bit Array that encodes the in-dictionary values |
| Secondary Table Metadata | .stab/.meta data | Stream | Json encoded secondary table metadata, e.g. `{ "format":"SimpleKV", "record_format":"range", "partitions":[ ["1",0,10000000], ["1",10000000,20000000], ["1",20000000,30000000] ], "compression":"NoCompression" }` |
| Secondary Table Partitions | .stab/<part-id> | Stream | A list of records |

#### Secondary Table Record

In the metadata, the field record_format indicates how the out-of-range value is represented, currently we should use “range” which indicates we use intervals. The logical layout is following

| Offset (bytes) | Name | Type | Note |
| --- | --- | --- | --- |
| 0..4 | left_pos | u32_le |  |
| 4..6 | size_enc | u16_le | length - 1 |
| 6..10 | value | i32_le |  |

For an inflated secondary table, it’s simply a stream of intervals.

Otherwise a deflated secondary table, each frame contains a metadata block

For an inflated secondary table, it's simply a stream of intervals. Otherwise a deflated secondary table, each frame contains a metadata block
| Offset (bytes) | Name | Type | Note |
| --- | --- | --- | --- |
| 0..4 | first_left_pos | u32_le |  |
| 4..? | Deflated records |  |  |

## References

1. Sasani, T. A. et al. Large, three-generation human families reveal post-zygotic mosaicism and variability in germline mutation accumulation. Elife 8, (2019).

2. Robinson, M. D., McCarthy, D. J. & Smyth, G. K. edgeR: a Bioconductor package for differential expression analysis of digital gene expression data. Bioinformatics 26, 139–140 (2010).

3. Anders, S. & Huber, W. Differential expression analysis for sequence count data. Genome Biol. 11, R106 (2010).

4. Pedersen, B. S., Collins, R. L., Talkowski, M. E. & Quinlan, A. R. Indexcov: fast coverage quality control for whole-genome sequencing. Gigascience 6, 1–6 (2017).

5. Kent, W. J., Zweig, A. S., Barber, G., Hinrichs, A. S. & Karolchik, D. BigWig and BigBed: enabling browsing of large distributed datasets. Bioinformatics 26, 2204–2207 (2010).

6. Koranne, S. Hierarchical Data Format 5: HDF5. Handbook of Open Source Tools 191–200 (2011) doi:10.1007/978-1-4419-7719-9_10.

7. ENCODE Project Consortium. An integrated encyclopedia of DNA elements in the human genome. Nature 489, 57–74 (2012).

8. Shao, Z., Reppy, J. H. & Appel, A. W. Unrolling lists. SIGPLAN Lisp Pointers VII, 185–195 (1994).

9. Li, H. et al. The Sequence Alignment/Map format and SAMtools. Bioinformatics 25, 2078–2079 (2009).

10. Pedersen, B. S. & Quinlan, A. R. Mosdepth: quick coverage calculation for genomes and exomes. Bioinformatics 34, 867–868 (2018).

11. Li, H. Toward better understanding of artifacts in variant calling from high-coverage samples. Bioinformatics 30, 2843–2851 (2014).

